# An optimized protocol for assay for transposase-accessible chromatin by sequencing (ATAC-seq) library preparation from adult *Drosophila melanogaster* neurons

**DOI:** 10.1101/2021.05.05.442838

**Authors:** Collin B. Merrill, Miguel A. Pabon, Austin B. Montgomery, Aylin R. Rodan, Adrian Rothenfluh

## Abstract

Assay for transposase-accessible chromatin by sequencing (ATAC-seq) is rapidly becoming the assay of choice to investigate chromatin-mediated gene regulation, largely because of low input requirements, a fast workflow, and the ability to interrogate the entire genome in an unbiased manner. Many studies using ATAC-seq use mammalian or human-derived tissues, and established protocols work well in these systems. However, ATAC-seq is not yet widely used in *Drosophila*. Vinegar flies present several advantages over mammalian systems that make them an excellent model for ATAC-seq studies, including abundant genetic tools that allow straightforward targeting, transgene expression, and genetic manipulation, and are not available in mammalian models. Because current ATAC-seq protocols are not optimized to use flies, we developed an optimized workflow that accounts for several complicating factors present in *Drosophila*. We examined parameters affecting nuclei isolation, including input size, freezing time, washing, and possible confounds from retinal pigments. Then, we optimized the enzymatic steps of library construction to account for the smaller *Drosophila* genome size. Finally, we used our optimized protocol to generate ATAC-seq libraries that meet ENCODE quality metrics. Our optimized protocol enables extensive ATAC-seq experiments in *Drosophila*, thereby leveraging the advantages of this powerful model system to understand chromatin-mediated gene regulation.

## INTRODUCTION

Assay for transposase-accessible chromatin by sequencing (ATAC-seq) is a relatively new technique that has advanced our understanding of chromatin-mediated gene regulation. The technique offers several advantages over other sequencing techniques that examine chromatin, such as micrococcal nuclease (MNase-seq) or formaldehyde-assisted isolation of regulatory elements (FAIRE-seq). These advantages include low input requirements, a fast workflow that does not require harsh chemicals, and genome-wide interrogation in an untargeted manner (Buenrostro et al., 2013). ATAC-seq is also robust to small sample sizes and has been adapted for single-cell applications (Cusanovich et al., 2015; Buenrostro et al., 2018).

The established ATAC-seq protocol was developed using peripheral blood samples (Buenrostro et al., 2013), and many studies using ATAC-seq focus on mammalian or human-derived tissues (Davie et al., 2015; Bysani et al., 2019; Liu et al., 2019). Recent studies used ATAC-seq to investigate other tissue types, including tissues from plants (Lu et al., 2017), zebrafish (Quillien et al., 2017), and livestock (Halstead et al., 2020). Some ATAC-seq studies using *Drosophila melanogaster* have been published, though many of these studies focus on embryogenesis and larvae (Cusanovich et al., 2018; Haines and Eisen, 2018; Bozek et al., 2019). One recent report performed ATAC-seq from central brains after fluorescent sorting of GFP-positive cell bodies (Brovkina et al., 2021). However, the physical dissection and dissociation of many brains can be labor-intensive and yield relatively few cells, in that case 6,000-10,000 cells (Brovkina et al., 2021). We were interested in developing a streamlined protocol that would allow efficient ATAC-seq library preparation, while taking advantage of the many genetic tools available for labeling specific *Drosophila* neurons (Jenett et al., 2012).

ATAC-seq library preparation has several parameters that can be optimized based on the input tissue. Many of these parameters are addressed in the original ATAC-seq protocol (Buenrostro et al., 2013). Additional studies optimized ATAC-seq library preparation from various tissue types in several storage conditions (Corces et al., 2017; Ho et al., 2018; Wang et al., 2018; Fujiwara et al., 2019). Most studies in *Drosophila* larvae treat the tissue similarly to mammalian tissues, only adding an additional washing step to clean the embryos or larvae before homogenization (Koenecke et al., 2016; Haines and Eisen, 2018). ATAC-seq library preparation using adult *Drosophila* poses challenges that are not present in other tissues. For example, the insect cuticle is a chitin structure that is difficult to process by simple homogenization. Another challenge is eye color, which is commonly used to identify the presence of transgenic insertions, and many transgenic fly lines contain the *mini-white* (*w*^*+mC*^) gene. In a *w*^*-*^ background, *mini-white* confers a yellow to red eye color, so eye color can range from white (*w*^*-*^) to red, depending on the genotype. Some *Drosophila* retinal pigments emit fluorescence (Franceschini et al., 1981; Miller et al., 1984) that can complicate fluorescent-activated nuclei sorting (FANS). Finally, the *Drosophila* genome is considerably smaller than mammalian genomes (Vinogradov, 2004; Canapa et al., 2016), so established ATAC-seq parameters may use higher-than-optimal enzyme concentrations and long incubation times that may lead to low-quality ATAC-seq libraries when prepared from fly tissues. Thus, optimizing each parameter for ATAC-seq library preparation using adult flies should be performed prior to sequencing. However, these experiments add considerable sequencing costs, putting the approach out-of-reach for many labs.

Identifying cell- and tissue-specific roles in organism function is a major goal of many current studies, which require specific and sensitive detection of target cells, especially for rare cell types. To facilitate these studies, we developed a protocol to generate ATAC-seq libraries from adult *Drosophila* neurons. Our goal was to optimize each variable in the library preparation process, with particular attention to challenges specific to *Drosophila* model systems. Our protocol provides guidelines for ATAC-seq library preparation using *Drosophila* and outlines the optimal parameters for each library preparation step. This protocol will accelerate studies of chromatin-mediated gene regulation in flies, thereby coupling the powerful genetic tools in *Drosophila* with high-throughput, in-depth examination of chromatin structure.

## MATERIALS AND METHODS

### Fly strains

The following fly lines were obtained from the Bloomington Drosophila Stock Center (Indiana University, Bloomington, IN, USA) and used in this study: *Canton S* (BL64349), *tub-Gal4* (BL5138), *elav*-*Gal4* (BL458), *nSyb-Gal4* (BL51635) *UAS-GFP-nls* (nuclear localization signal; BL4775), *UAS-Stinger* (a super-bright GFP-nls variant; BL84277), *TH*-*Gal4* (BL8848), *Cha-Gal4*.*19B* (BL6798), and *vGAT*-*Gal4* (BL58980). *w* Berlin* flies have been in the lab for many generations (Wolf et al., 2002) and were originally a gift from Martin Heisenberg. Flies were reared in bottles containing standard cornmeal agar and grown at 25 °C with 70% relative humidity and a 12-h light/dark cycle.

### Nuclei isolation

All equipment and buffers were pre-chilled to be ice-cold. All plastic and glassware used for nuclei isolation was pre-treated with 1% bovine serum albumin (BSA) in phosphate-buffered saline (PBS). Flies were collected into empty bottles and frozen at -80 °C. Then, the flies were placed in pre-chilled (in a -80 °C freezer) sieves and agitated for 1 min to separate the heads from the bodies. The fly heads were added to lysis buffer [10 mM Tris-HCl pH 7.5, 10 mM NaCl, 3 mM MgCl_2_, 0.1% Tween-20, 0.1% Nonidet P40 substitute, 0.01% digitonin (ThermoFisher Scientific, Waltham, MA, USA), and 1% bovine serum albumin] in a Dounce homogenizer. The tissue was homogenized with the A pestle until the resistance disappeared. The crude homogenate was passed through a 40 μm filter and homogenized with 15 slow strokes with the B pestle. The crude nuclei were washed with wash buffer (10 mM Tris-HCl, 10 mM NaCl, 3 mM MgCl_2_, 1% BSA, and 0.1% Tween-20) and centrifuged at 500 *g* for 10 minutes per wash. The washed nuclei were resuspended in 1 mL wash buffer with 3 mM DAPI for evaluation and sorting.

### Fluorescence-activated nuclei sorting (FANS)

Nuclei were evaluated and sorted with a BD FACS Aria flow cytometer (BD Biosciences, San Jose, CA, USA) operated by the University of Utah Flow Cytometry Core facility. Nuclei collected from *w* Berlin* flies (no GFP) were stained with 3 μM DAPI and used as the GFP-negative control to set the sorting gates. GFP-positive nuclei were collected into 500 μL wash buffer and stored on ice until use for ATAC-seq library prep. Nuclei counts were determined using an internal control bead population (Spherotech, Lake Forest, IL, USA) at 10^6^ beads/mL. Briefly, 20 uL of well-mixed beads were added to 200 uL of nuclei suspension and gently mixed. Data was recorded for the bead and cell mixture until at least 10,000 singlet beads were collected. Nuclei were identified based on the forward scatter signal and DAPI intensity. The number of nuclei was calculated as follows: number of nuclei recorded × 10^5^/number of beads recorded = number of nuclei/mL.

### DNA tagmentation, amplification, and purification

Purified nuclei were centrifuged for 10 min at 500 *g* at 4 °C. The pellet was mixed with 22.5 μL 2X Tn5 tagmentation buffer [20 mM Tris-HCl pH 7.6, 10 mM MgCl_2_, and 20% dimethyl formamide (Sigma Aldrich, St. Louis, MO, USA) in sterile water], 16.5 μL 1X PBS, 0.5 μL Tween-20, 5.5 μL water, and Tn5 enzyme (Illumina, Inc., San Diego, CA, USA). The mixture was incubated at 37 °C for various times and the DNA was purified using a MinElute PCR purification kit (Qiagen, Germantown, MD, USA). Then, the purified DNA was mixed with CD index primers (Illumina) and Phusion High Fidelity PCR MasterMix (New England Biolabs, Ipswich, MA, USA) and amplified by 1 cycle of 72 °C for 5 min and 98 °C for 30 sec, followed by 5 cycles of 98 °C for 10 sec, 63 °C for 30 sec, and 72°C for 1 min in a C100 thermocycler (BioRad, Hercules, CA, USA). 5 μL of this reaction was removed and used for a qPCR side reaction. The aliquot was mixed with the same CD index primers and SsoFast EvaGreen Supermix (BioRad) and amplified for 1 cycle of 98 °C for 30 sec and 40 cycles of 98 °C for 10 sec, 63 °C for 30 sec, and 72 °C for 1 min in an Applied Biosystems 7900HT qPCR instrument (ThermoFisher Scientific). Total fluorescence was calculated by subtracting the fluorescence from first five PCR cycles from the maximum fluorescence of each sample. The cycle number corresponding to a portion of the total fluorescence was determined (∼8-10 cycles) and the DNA was amplified for this number of additional cycles. After amplification, the DNA was purified with 0.5X and 1.1X AMPure XP beads (Beckman Coulter, Indianapolis, IN, USA) to remove primer-dimers and high-molecular weight DNA (> 5000 bp). The purified DNA was stored at -20 °C until further analysis.

### ATAC-seq library quality assessment

ATAC-seq library quality was assessed using an Agilent 2200 TapeStation and High Sensitivity D5000 ScreenTape assays (Agilent Technologies, Inc., Santa Clara, CA, USA) at the Huntsman Cancer Institute High Throughput Genomics core facility or the Genome Technology Access Center of the McDonnell Genome Institute at the Washington University School of Medicine (St. Louis, MO, USA).

### Next generation sequencing and analysis

ATAC-seq libraries constructed using nuclei isolated from dopaminergic and GABAergic neurons were sequenced on a Novaseq 6000 instrument (Illumina) using 50-bp paired-end reads. Fastq files were quality checked using FastQC (v 0.11.9). Adapter sequences were removed with CutAdapt (v3.4). The sequenced reads were aligned to the dmel_r6.26 genome assembly using Bowtie2 (v2.4.2; Langmead and Salzberg, 2012). Aligned reads were sorted with Samtools (v1.12; Li et al., 2009) and deduplicated with Picard (v2.23) using the MarkDuplicates command. Aligned, filtered, and deduplicated reads were used to call peaks using MACS2 software (v2.2.5; Zhang et al., 2008). Then, featureCounts (v2.6.4; Liao et al., 2019) was used to generate the count matrix. Differential accessibility analysis was performed using DESeq2 (v1.32.0; Love et al., 2014). Insert size distributions were calculated using Picard. Peaks were visualized using the IGV browser (v2.7.2).

### Statistical analysis

The data are represented as means ± standard deviation. Comparisons between homogenizer size and fly volume were analyzed by two-way ANOVA with Bonferroni ‘s multiple comparison tests. Freezing time was analyzed by one-way ANOVA using Tukey’s multiple comparison tests. Eye color experiments, % read alignment, and fractions of reads in peaks (FRiP) were analyzed using Student’s t-tests. Gal4-UAS and washing experiments were analyzed by simple linear regression, and analysis of covariance (ANCOVA). All statistical analyses were performed using GraphPad Prism 9 software (San Diego, CA, USA).

## RESULTS

### Sample volumes, homogenization, and freezing time

Our goal was to identify the optimal conditions for nuclei isolation, purification, and ATAC-seq library preparation using adult *Drosophila*. We first asked if the number of flies used for head isolation and the size of the Dounce homogenizer play a role in the number of extracted nuclei that are labeled with DAPI. Using a larger volume of flies increased the number of extracted nuclei (2-way ANOVA, F(1,8) = 15.61, *p =* 0.0042). The size of the Dounce homogenizer used for extraction did not significantly affect the total extracted nuclei if analyzed by 2-way ANOVA (F(1,8) = 1.984, *p* = 0.1966; Fig. 1A), however, with 0.5 mL of flies, more nuclei were isolated with the 7 mL homogenizer (significant by isolated t-test). Thus, we recommend using the larger, 7 mL Dounce homogenizer to maximize nuclei extraction.

**Figure 1.**
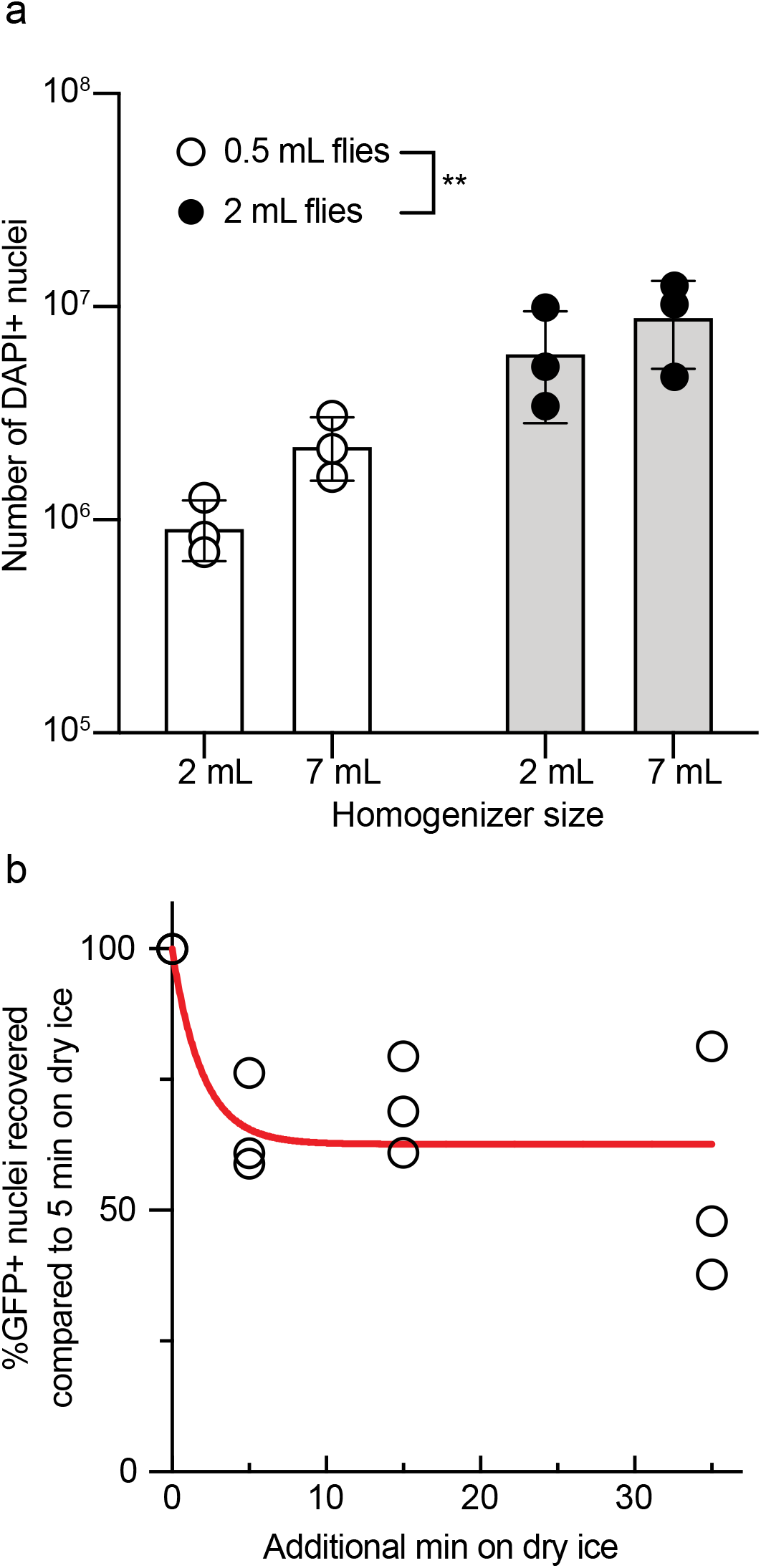
Sample volume and freezing time affect nuclei yield. A) The number of flies used to collect heads prior to nuclei isolation increases the number of extracted nuclei, but the homogenizer size does not. The indicated volumes of *w* Berlin* flies were frozen, the heads shaken off in a sieve, the nuclei were extracted and stained with DAPI, and were then counted in a flow cytometer. Data represent the mean ± SD of three independent experiments. B) Freezing time longer than 5 min decreases GFP detection by flow cytometry. Flies with pan-neuronal GFP, *nSyb-Gal4 UAS-GFP-nls* were counted at the low gate. Data represent three biological replicates. The data in A) were analyzed by two-way ANOVA and the data in B) were analyzed by nonlinear regression using one-phase exponential decay, K = 0.51 min^-1^, goodness of fit *R*^*2*^ = 0.64. \*\**p* < 0.01.

One advantage to using *Drosophila* is the availability of sophisticated genetic tools that allows the easy identification of target cells, which can then be isolated by flow cytometry. In our nuclei isolation procedure, *nSyb-Gal4* flies expressing pan-neuronal GFP-nls are frozen at -80 °C for 5 min and passed through a fly sieve to isolate the tissue of interest (bodies vs. heads). We hypothesized that extended freezing at -80 °C could reduce GFP fluorescence, so we performed a time-course experiment to examine whether freezing time influences GFP fluorescence. Five min is about the minimum time per sample in which flies can be frozen at -80 °C and homogenized when handling multiple samples. We observed that GFP fluorescence decreased quickly with additional time on dry ice but reached a plateau at ∼60% fluorescence in the time analyzed (Fig. 1B, one-phase exponential decay, K = 0.51 min^-1^, goodness of fit *R*^*2*^ = 0.64). Therefore, we suggest that flies should only be frozen briefly (5 min or less) at -80 °C when used for nuclei isolation and ATAC-seq library preparation.

### Flow cytometer specificity and sensitivity

Sorting GFP+ nuclei using a flow cytometer is dependent on the gating scheme for the detection of green fluorescent signal. Detection schemes for stains usually come at a low gate (less signal required) and a high gate (more signal required). To determine the sorting sensitivity at these gates, we ubiquitously expressed a brighter version of nuclear GFP, *UAS-Stinger*, using a strong *Tubulin-Gal4* driver. At the low gate, we recovered ∼80% of nuclei from ∼100,000 DAPI+ nuclei (Fig. 2A), while we recovered ∼12% of nuclei with the high gate (Fig. 2B). An independent experiment showed similar percentages. The data from these experiments suggest that the low gate has 6-7 fold better sensitivity than the high gate.

**Figure 2.**
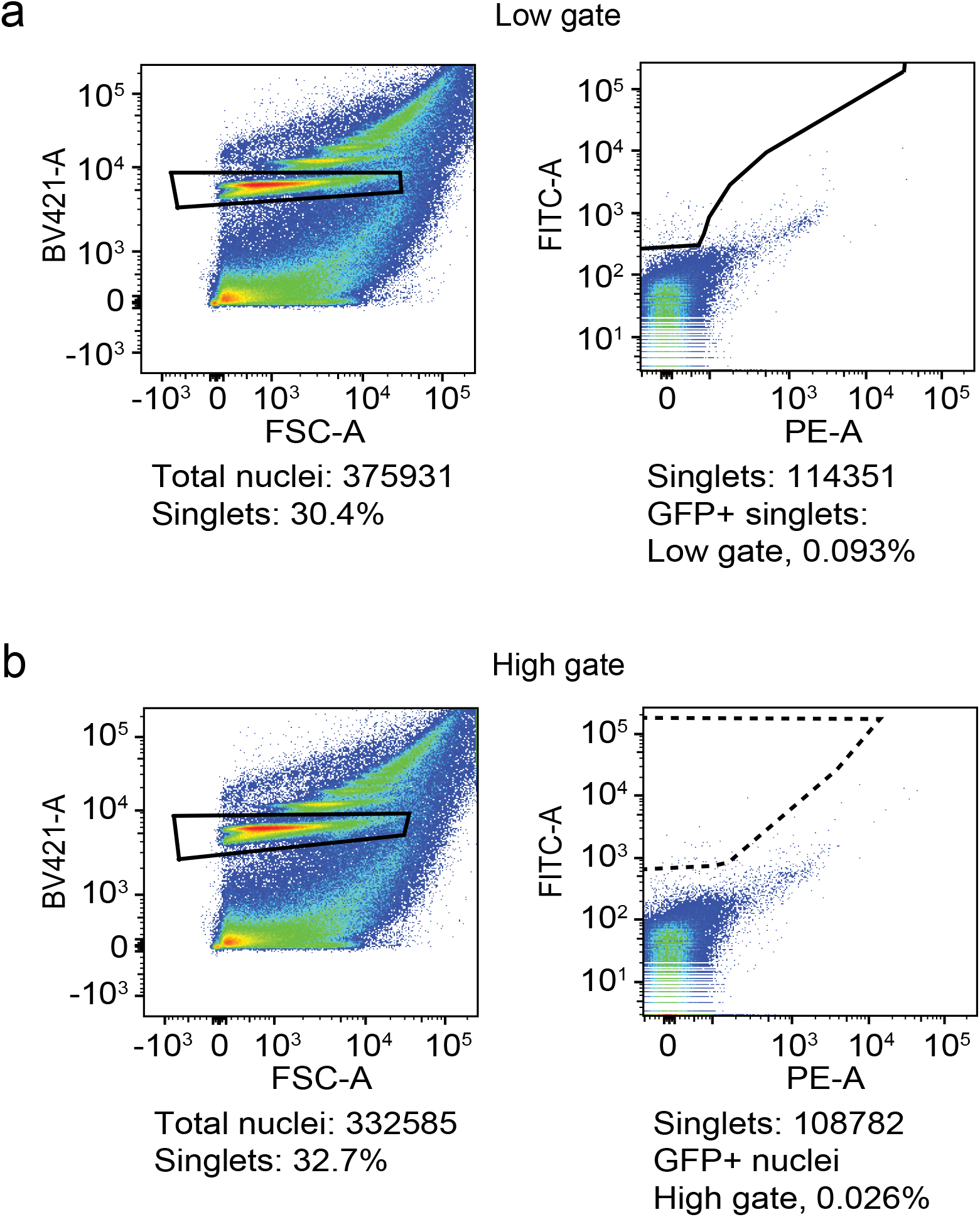
Flow cytometry shows high specificity and few false positives. Nuclei were isolated from *w* Berlin* heads (GFP-negative) and analyzed by flow cytometry. A) Representative dot plots of nuclei analyzed using the low gate. Nuclei were identified by DAPI staining (Y-axis) and forward light scatter (X-axis, left). The box indicates the sorted DAPI+ singlet nuclei, which were then analyzed for green fluorescence (Y-axis) and the red PE-A channel (X-axis, right), which allowed the best separation of GFP populations, and sorted at the low gate (solid trapezoid box). 0.093% of input singlets were GFP false-positives (and 0.024% when applying the high gate in this sort). B) Representative dot plots of nuclei analyzed using the high gate. Nuclei were isolated as in A), shown on the left, and then sorted for green fluorescence at the high gate (stippled trapezoid box, right). 0.026% of input singlets were GFP false-positives (and 0.10% when applying the low gate in this sort).

To examine the sorting specificity, we evaluated the number of sorted ‘GFP+ nuclei’ in *w* Berlin* flies, which are all GFP-negative. Sorting at the low gate yielded 0.093% ‘GFP+’ nuclei (Fig. 3A), while at the high gate, 0.026% of nuclei were ‘GFP+’ (Fig. 3B). An independent experiment gave the same results. Together, these data suggest that the high gate has 4-fold higher specificity and is better at calling true positives. As expected, there is a tradeoff between sensitivity and specificity when using low and high GFP+ sorting gates.

**Figure 3.**
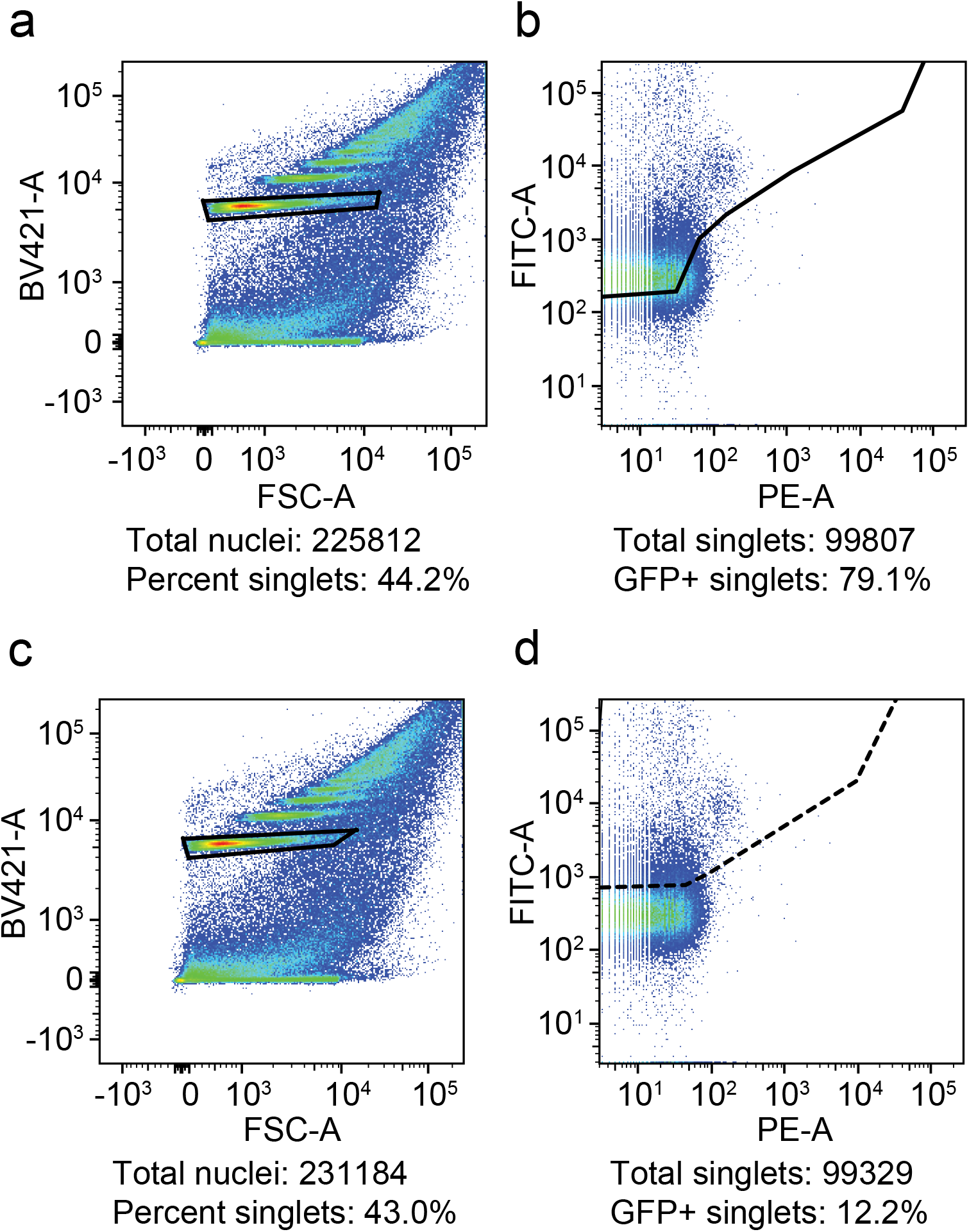
Differential sensitivity at the low and high sorting gate. *UAS-Stinger*, encoding a super-bright nuclear GFP, expression was driven using ubiquitous *Tubulin*-*Gal4* (all nuclei are GFP+). Nuclei from heads were isolated as described in Fig. 2 (left sides) and analyzed by flow cytometry using low- and high-stringency gating (right). A) 79% of nuclei were called as GFP+ at the low gate (solid trapezoid). B) 12% of nuclei were called as GFP+ at the high gate (stippled trapezoid, right).

### Drosophila-specific parameters affect nuclei isolation

Using the high sorting gate, 1 in ∼4000 nuclei are selected as false-positive. At face value, this is good specificity. However, assuming that a *Drosophila* head contains >200,000 nuclei, >50 of those will be sorted as false positives at the high gate (and >200 at the low gate). These numbers would significantly contaminate the nuclei sorted from a genotype where Gal4 is expressed in a subset of 100s of neurons. The high gate will produce fewer false positives, thus increasing the specificity for the Gal4-expression pattern, but the high gate will also reduce the sensitivity and recovery of those Gal4+ neurons. Therefore, we asked whether we could increase the recovery via genetic means by increasing the number of *Gal4* or *UAS-GFP-nls* transgenic constructs to boost GFP expression. We used *ics-Gal4 UAS-GFP-nls* flies and crossed them to themselves, *ics-Gal4, UAS-GFP-nls*, or *w* Berlin* to generate flies with 4, 3, 3, or 2 copies of *Gal4/UAS* insertions, respectively. *ics-Gal4* expression includes many neurons and the expression pattern is not different between *ics-Gal4/+* hetero- and *ics-Gal4* homozygotes. (Ojelade et al., 2015). Expanding the copies of *Gal4/UAS* insertions increased the number of detected GFP+ nuclei (linear regression, slope ≠ 0, *p* = 0.0008, low gate; *p* = 0.001, high gate, Fig. 4A). At both gates, the recovery with 4 transgenes was about 20-fold higher than recovery with 2 transgenes. We therefore recommend that for Gal4-lines with sparse expression patterns, the amount of GFP produced should be increased genetically.

**Figure 4.**
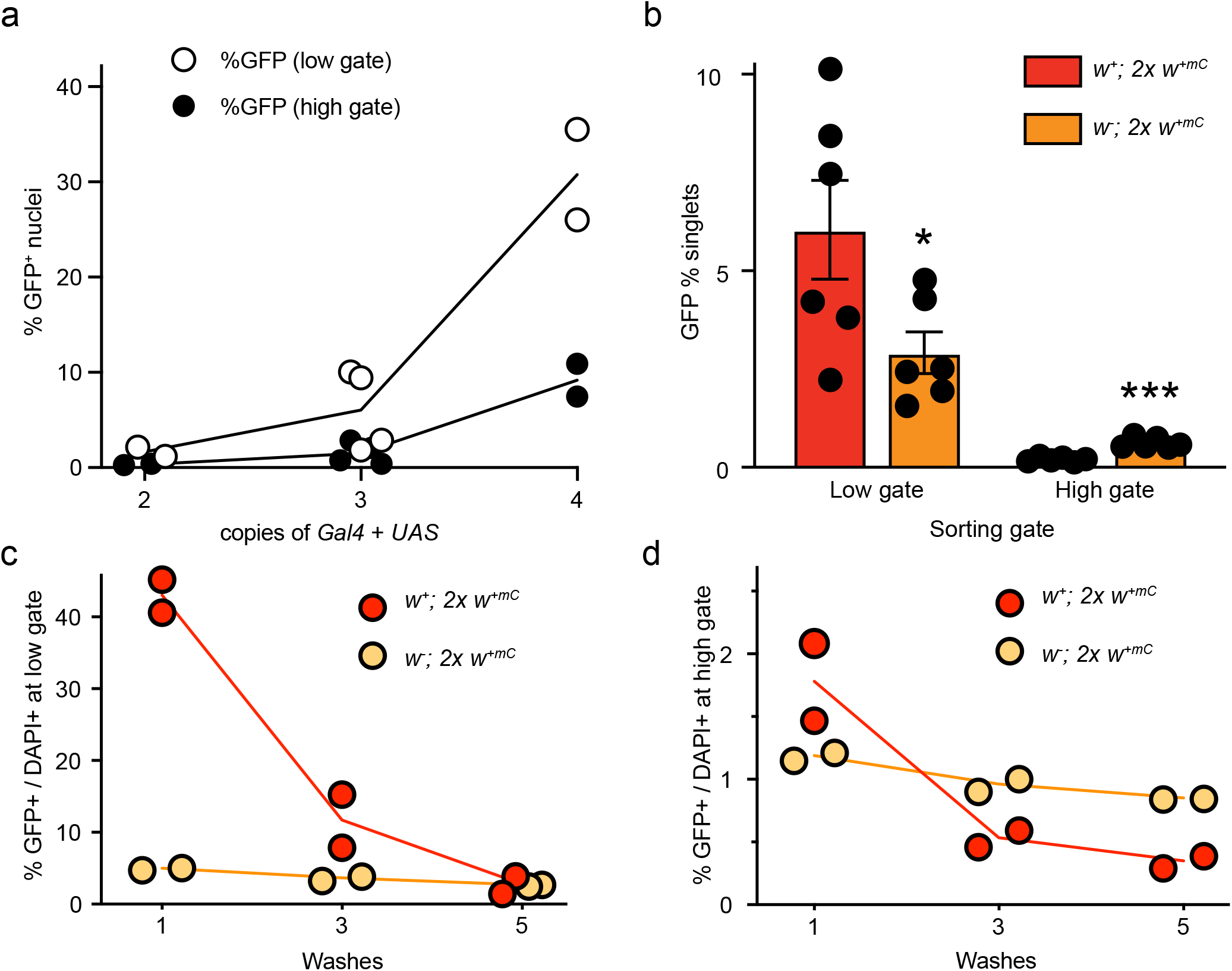
Physical parameters affecting ATAC-seq library preparation from *Drosophila* nuclei. ChangesToFig. A) GFP+ nuclei from flies expressing multiple copies of Gal4 + UAS (*ics-Gal4 UAS-GFP-nls*) were isolated and analyzed by flow cytometry using low- and high-stringency gating. B) Nuclei from cholinergic neurons (heads of *Cha-Gal4 UAS-GFP-nls*) red-eyed (*w*^*+*^ background) and orange-eyed flies (*w*^*–*^ background) were isolated and analyzed by flow cytometry using low- and high-stringency gating. Error bars represent the mean ± SD of six biological replicates and were analyzed by Student’s t-tests. C-D) Nuclei from the same flies used in B) were washed multiple times after isolation and before analysis using low- (C) and high- (D) stringency gating. The data was analyzed using linear regression. All regression lines have slopes ≠ 0, *p* < 0.02.

Most *Drosophila* transgenic insertions are marked with a miniature copy of the *white*^*+*^ gene, *w*^*+mC*^. The presence of such a transgene will confer anywhere from pale yellow to red-colored eyes. In addition, the allele *w*^*+mC*^ is dosage-sensitive, *i*.*e*. the more transgenes in the fly labeled with *w*^*+mC*^, the closer to wild-type red the eye color will be. Since we suggested above that increasing the copy of *Gal4* and/or *UAS* transgenes will increase the recovery of GFP+ nuclei, we next wanted to determine if red-colored eye pigment interferes with sorting GFP+ nuclei. We crossed *w*^*-*^*;Cha-Gal4 UAS-GFP-nls* males to white-eyed (*w*^*-*^), or to red-eyed (*w*^*+*^) virgins and then collected the male progeny. Both of these male genotypes contain 2 *w*^*+mC*^ alleles from the transgenes, but one group is red-eyed (*w*^*+*^) and the other is orange-eyed (in a *w*^*-*^ background). At the low gate, we recovered almost twice as many ‘GFP+ nuclei’ (Fig. 4B), suggesting that red eye pigment from *w*^+^ contributes to the false positive rate. We hypothesized that increasing the number of washes during nuclei isolation would mitigate the false positive rate caused by red eye pigments. Washing the crude nuclei extracts one, three-, or five-times dose-dependently reduced the GFP+/DAPI+ ratio in red-eyed and orange-eyed flies. For the red-eyed flies, we observed the steepest decrease between one and three washes, and with three washes, there was no longer a difference in the GFP+/DAPI+ ratio between red- and orange-eye flies (Fig. 4C,D). We therefore recommend at least three washes, especially when sorting with the low-stringency gate and as more *w*^*+mC*^-marked transgenes are used, since the eye color will approach wild-type red.

### Reaction conditions affect library quality

Published ATAC-seq guidelines are based on mammalian genomes, which are approximately 10-fold larger than the *Drosophila* genome (Vinogradov, 2004; Canapa et al., 2016). Therefore, the parameters used for the Tn5 transposition reaction (incubation time and enzyme concentration) in mammalian cells may not be appropriate for fly nuclei. To determine the optimal Tn5 reaction conditions, we sorted GFP+ nuclei from fly heads and constructed ATAC-seq libraries. We evaluated the library quality using the guidelines recommended by Buenrostro et al. (2015). We first examined how reaction time affects the fragment distribution by sorting pan-neuronal GFP+ nuclei from *nSyb-Gal4 UAS-GFP-nls* fly heads. Then, we prepared ATAC-seq libraries using 1X Tn5 enzyme and sample incubation for 5, 23, and 60 min (Fig. 5A). A low reaction time decreased the nucleosome-free peak and increased the proportion of larger fragments up to 1000 bp, suggesting that di-, tri- and multi-nucleosome fragments are over-represented. Incubation for 23 min provided a fragment distribution with more nucleosome-free and fewer mono- and di-nucleosome fragments, while increasing the reaction time to 60 min biased the library toward smaller fragment sizes with relatively little evidence of larger fragments. Thus, for library preparation using *Drosophila* nuclei, 23 minutes is the optimal reaction time. We next examined how the Tn5 concentration affects libraries by collecting GFP+ nuclei from *tubulin-Gal4 UAS-stinger-nls* flies and tagmenting the DNA using 0.1X, 0.3X, 1X, and 3X Tn5 (compared to manufacturer recommendation; Fig. 5A). 1X Tn5 showed the largest peak around 200 bp, with additional skewing towards larger fragments. Reacting DNA using 3X Tn5 reduced both skewness toward larger fragments and the number of nucleosome-free fragments due to over-digestion of DNA [which was size-filtered (150∼2000 bp) before analysis, Fig. 5B]. Reducing the Tn5 concentration to 0.1X showed skewness to higher fragments, but overall fewer amplified fragments and reduced overall recovery. We therefore recommend 1X Tn5 as the best balance between recovery of amplified fragments and avoiding over-digestion. Finally, we examined how library amplification affects quality. Like other sequencing library types, ATAC-seq libraries must be amplified to obtain sufficient DNA concentrations for sequencing. Buenrostro et al. (2013, 2015) suggested that ATAC-seq libraries should be amplified to 33% of maximal fluorescence obtained in a qPCR pre-reaction to ensure that sufficient material is present for sequencing, without the introduction of GC and size bias. We hypothesized that the smaller genome size in *Drosophila* would require more PCR cycles than libraries derived from mammalian nuclei. We tagmented DNA from pan-neuronal GFP+ nuclei from *nSyb-Gal4 UAS-GFP-nls* fly heads with 1X Tn5 and amplified the tagmented DNA to 25% (8 cycles), 33% (9 cycles), and 50% (10 cycles) of total fluorescence in a qPCR side reaction (40 cycles) to determine the number of PCR cycles needed for library amplification before sequencing (Fig. 5C). The library that was amplified to 33% total fluorescence had the smallest amount of nucleosome-free fragments and a large proportion of ∼800-1200 bp fragments, which may correspond to a PCR bubble that arose from depleted PCR reagents, most likely primers (Kanagawa, 2003). In contrast, the library amplified to 25% total fluorescence showed a broad nucleosome-free peak and an even distribution of higher molecular weight fragments, while the library amplified to 50% total fluorescence showed a tall nucleosome-free peak that suggests a bias toward low molecular weight amplification. These results suggest that ATAC-seq libraries derived from *Drosophila* nuclei should be amplified to 25% of total qPCR fluorescence, though 33%-amplified libraries may also be suitable for sequencing after reconditioning PCR (Thompson et al., 2002).

**Figure 5.**
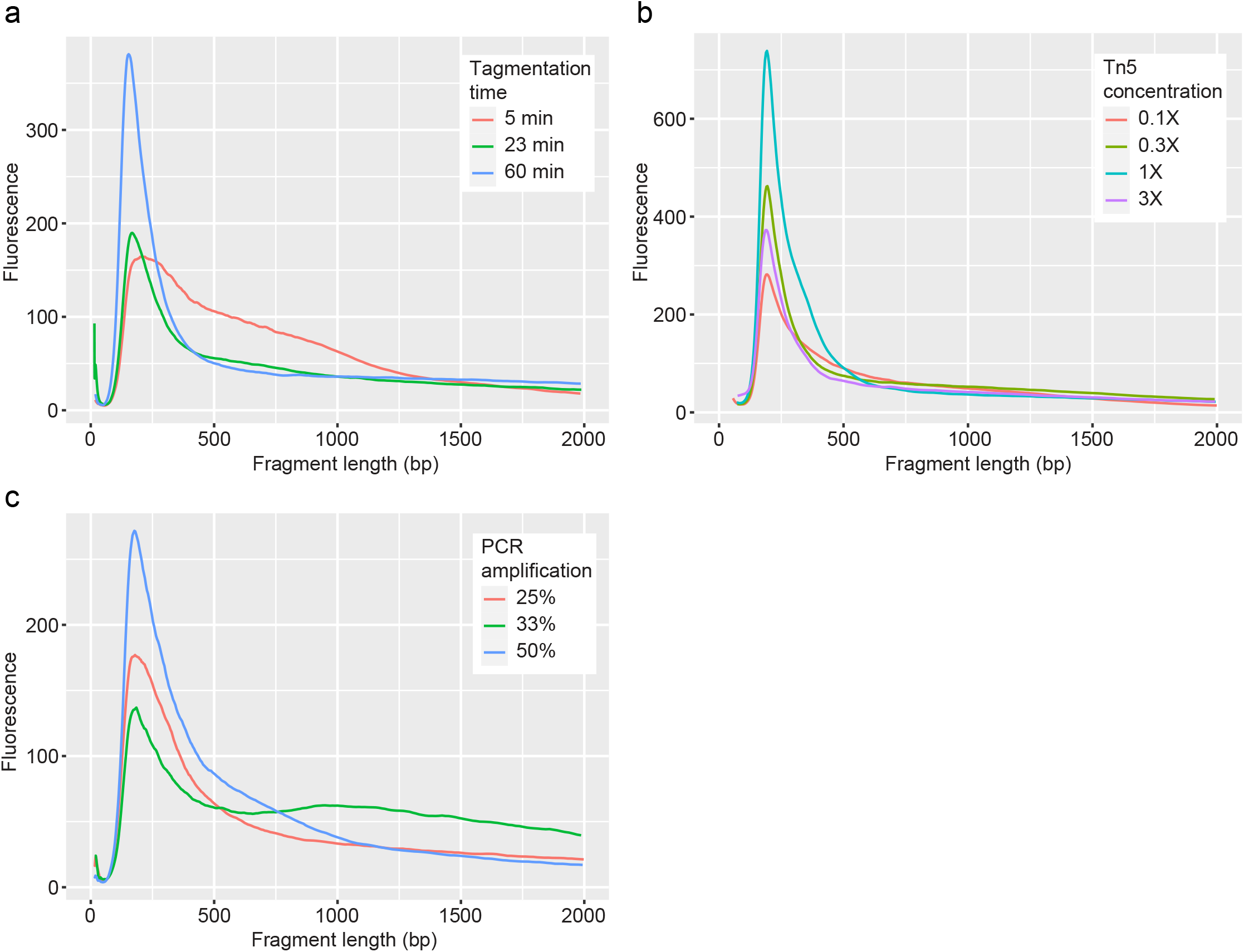
Optimal library construction parameters. GFP expression was driven using various fly genotypes. Then, nuclei were collected by flow cytometry using high-stringency gating. The isolated nuclei were used in ATAC-seq library construction, with variable preparation conditions as indicated. After amplification, each library was purified using AMPure XP beads, which remove low- and high-molecular weight fragments. The fragment sizes and relative fluorescence from Tapestation data are shown. A) Tn5 reaction time; B) Tn5 concentration; and C) post-Tn5 PCR amplification. GFP expression in A) was driven using *nSyb-Gal4 UAS-GFP-nls*; GFP expression in B) was driven using *Tubulin-Gal4 UAS-Stinger*; and GFP expression in C) was driven using *nSyb-Gal4 UAS-GFP-nls*.

### Optimized reaction conditions generate ATAC-seq libraries conforming to ENCODE standards

Finally, we combined our optimized parameters to generate and sequence ATAC-seq libraries. We drove nuclear GFP expression in dopaminergic and GABAergic neurons and isolated the nuclei after freezing approximately 1 mL (300-400) flies at -80 °C for 5 min. The crude nuclei extracts were washed three times and sorted by flow cytometry using a high-stringency gate. We collected 65,000 nuclei per sample for each neuron type, which were tagmented for 23 min using 1X Tn5 (from an Illumina Nextera kit). The resulting libraries were amplified to 25% qPCR fluorescence (8 cycles) and analyzed using 50-bp paired end sequencing. We then examined the percent alignment to the genome, the fraction of reads in peaks (FRiP), and fragment length distribution of the sequenced libraries. Both libraries showed high alignment rates, with 79% and 94% alignment in dopaminergic and GABAergic libraries, respectively (Fig. 6A). Dopamine neuron libraries had 0.37 FRiP, while GABA neuron libraries had 0.40 FRiP (Fig. 6B), which were both higher than the FRiP values (>0.3) recommended by the ENCODE consortium for ATAC-seq libraries (Davis et al., 2018). Finally, each of these libraries showed the expected nucleosomal banding pattern (Fig. 6C). Together, these metrics indicate that we generated high-quality libraries and that our protocol returns high-quality sequencing data. To test this hypothesis, we examined ATAC-seq peaks in regions associated with classical and neuron markers identified by single-cell RNA-seq (Davie et al., 2018). We observed differential peaks, indicating increased accessibility, in *ple, DAT*, and *hth*, which encode tyrosine hydroxylase, the rate-limiting dopamine-synthesis enzyme, the dopamine transporter, and homothorax, a homeobox transcription factor, respectively. These peaks were significantly more open in dopamine neurons than in GABA neurons (peaks indicated by asterisks in Fig. 7A, from left to right; *ple*: *p =* 3.32e-65 and 5.01e-53; *DAT*: *p* = 1.03e-05 and 1.04e-05; *hth*: *p* = 0.009, 1.69e-36, 1.14e-21, and 0.0008). Peaks associated with dopamine-neuron specific genes were much smaller or absent in GABA neurons (Fig. 7). The peaks corresponding to the GABAergic markers include *Gad1*, which encodes glutamic acid decarboxylase, which synthesizes GABA, *Lim3*, and *CG14989*. Peaks associated with GABA marker genes were significantly more open in GABA neurons than in dopamine neurons (peaks indicated by asterisks in Fig. 7B, from left to right; *GAD1*: *p* = 0.0005, 4.42e-08, 0.04, 0.01, and 6.6e-08; *Lim3*: *p* = 1.39e-05 and 0.001; *CG14989*; *p* = 0.1). These peaks were much smaller or absent in dopaminergic neurons. These results confirm that our nuclei isolation and library generation protocol yields high quality data, including differential chromatin accessibility in known cell-type markers (Davie et al., 2018).

**Figure 6.**
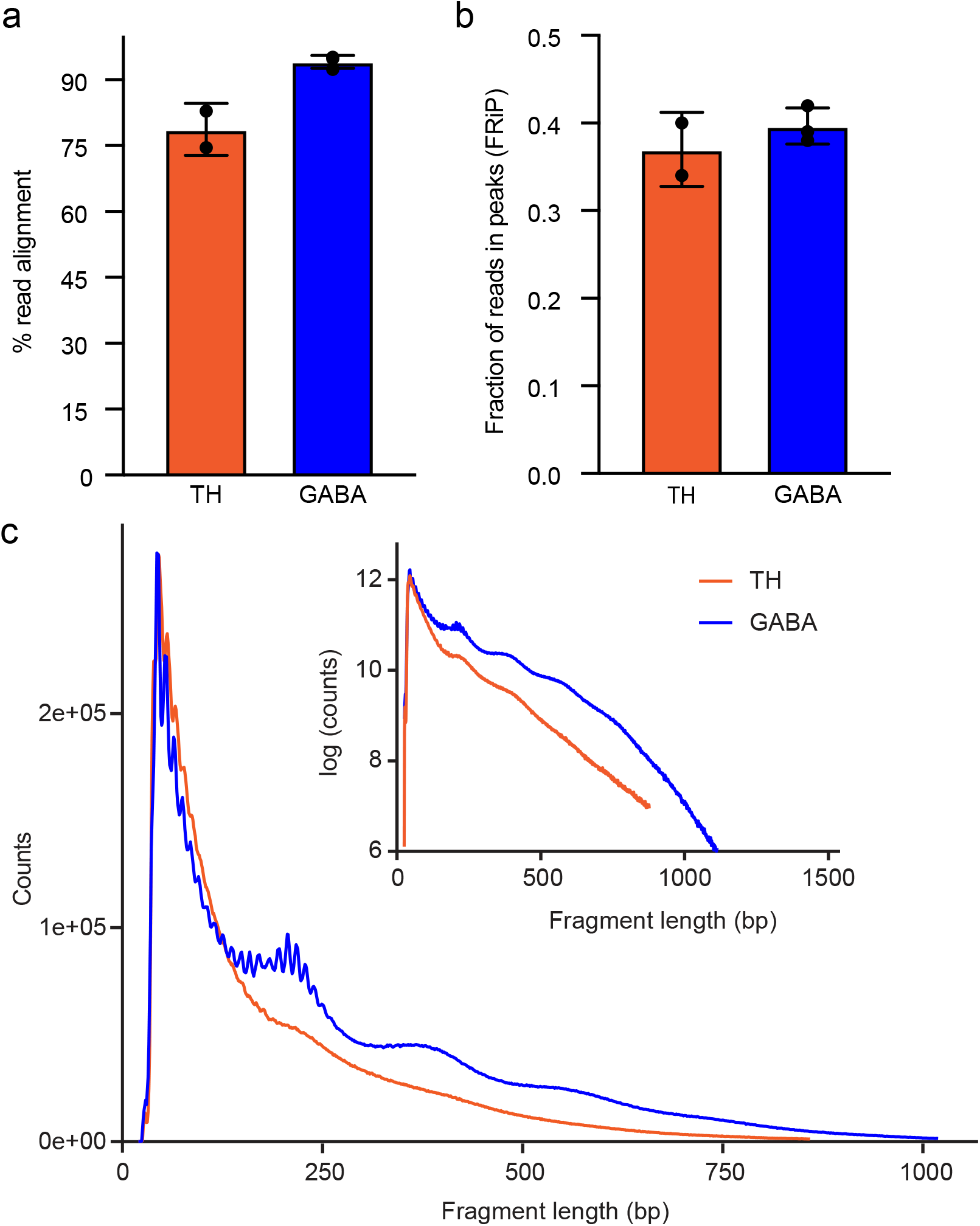
Optimized ATAC-seq parameters generate consistent libraries. Representative traces from ATAC-seq libraries prepared from dopaminergic and GABAergic nuclei isolated from adult *Drosophila* heads. The libraries were prepared using nuclear GFP+ expression driven by *TH*-*Gal4* (dopamine) and *vGAT*-*Gal4* (GABA). Flies were frozen at -80 °C for 5 min per replicate (300-400 flies per replicate; 3 replicates per genotype), the heads were collected, and the nuclei were isolated by flow cytometry using high-stringency gating. The target DNA was incubated for 23 minutes with 1X Tn5, and the libraries were PCR-amplified for 8 cycles, which corresponded to 25% total fluorescence from the qPCR side reaction. Amplified libraries were purified using AMPure XP beads and sequenced using a Novaseq 6000 instrument using 50 bp paired-end reads. A) Fraction of reads aligned to the nuclear genome. B) Fraction of reads in peaks (FRiP) after alignment, filtering, and peak calling. C) Insert size distribution showing the expected nucleosomal periodicity. Inset: Log-scaled insert size distribution. In A) and B), the data represent the mean ± SD of three replicates per neuron type. Statistical differences were analyzed using Mann-Whitney U tests.

**Figure 7.**
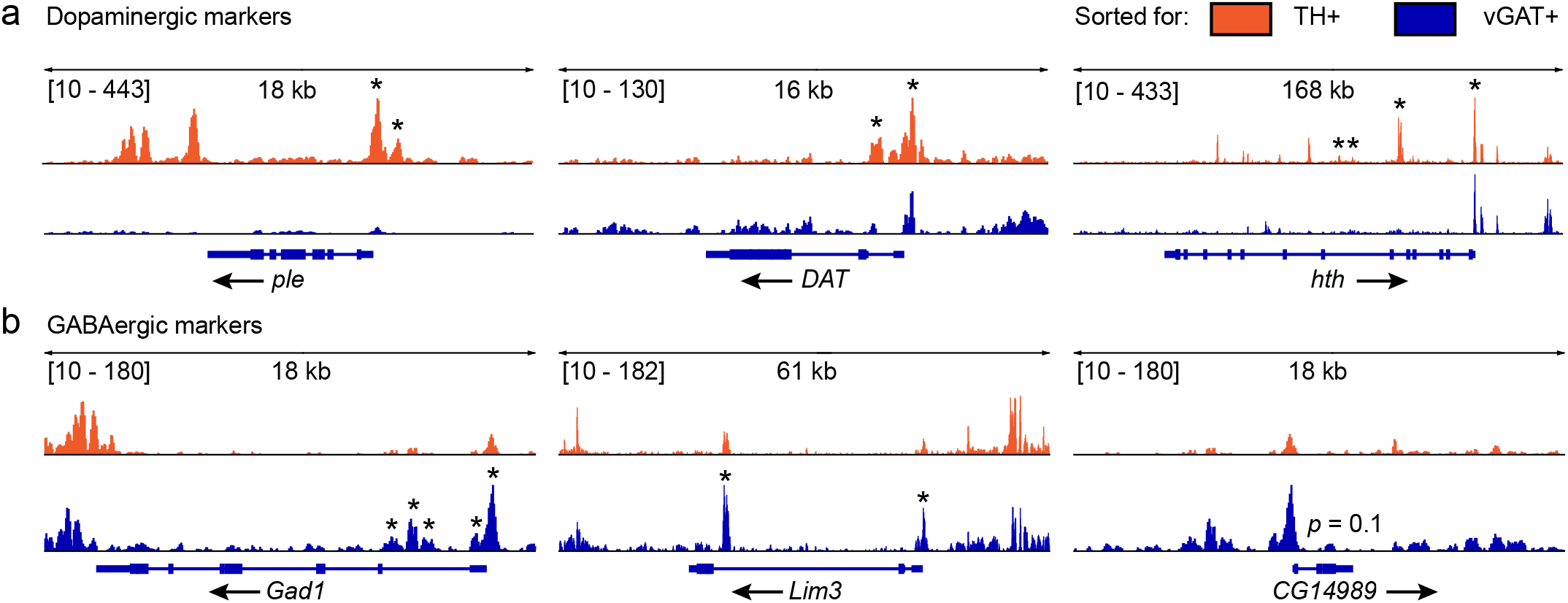
Optimized ATAC-seq libraries return expected cellular markers. ATAC-seq peaks in marker genes for dopaminergic and GABAergic neurons. ATAC-seq reads from dopamine neurons (tyrosine hydroxylase: TH+) are shown in orange and reads from GABA (vesicular GABA transporter: vGAT+) neurons are shown in blue. The genes and gene structures are shown below the ATAC-seq peaks. A) Peaks in chromatin regions that are more open in dopamine neurons. *ple* (encoding tyrosine hydroxylase; *p* = 2.43e-68) and *DAT* (dopamine transporter; *p* = 9.64e-7) are classical dopamine neuron markers. *hth* (homothorax; *p* = 1.16e-52) is a newly described dopamine neuron marker. B) Chromatin regions that are more open in GABA neurons. *Gad1* (glutamic acid decarboxylase 1; *p =* 6.64e-8) is a classical GABAergic neuron marker, while *Lim3* (Lim3; *p* < 0.0002) and *CG14989* (uncharacterized protein; *p* = 0.1) were identified as additional GABA neuron markers. *CG14989, Lim3*, and *hth* were described as specific markers using single-cell RNA sequencing (Davie et al., 2018). The boxes in the gene structures indicate exons and the arrows indicate the transcription direction.

## DISCUSSION

ATAC-seq is a powerful way to interrogate the chromatin state within tissues of interest. Critically, the technique provides key advantages over previous methods, such as ChIP-seq, MNase-seq, or FAIRE-seq, including low input requirements (50,000-60,000 nuclei vs. 1-10 million nuclei for MNase- or FAIRE-seq) (Kidder et al., 2011; Simon et al., 2012; McKay and Lieb, 2013), less hands-on time, no harsh chemicals such as paraformaldehyde, and no antibody optimization or complicated pull-down methods. Thus, ATAC-seq is quickly becoming the assay of choice for chromatin studies.

Most published studies using ATAC-seq focus on either tumor tissue or white blood cells, though recent studies have investigated the brain and other tissues in mice and humans (Liu et al., 2019; Rocks et al., 2021). The initial study describing ATAC-seq was performed in mammalian tissue (Buenrostro et al., 2013), and published protocols have been optimized for use in these tissues (Corces et al., 2017; Buenrostro et al., 2018). *Drosophila melanogaster* is a powerful model organism that is routinely used to study development and human disease models. Because we are interested in *Drosophila* neurons from the adult brain, we first determined some basic parameters for nuclei isolation. We found that freezing flies for ≥ 5 min and shaking off the heads in a sieve to enrich for brain tissue did not impede sorting GFP+ nuclei, but freezing flies for a longer duration did reduce GFP+ nuclei recovery. Using adult *Drosophila* – heads or whole flies – for ATAC-seq experiments presents some complicating factors, such as the exoskeleton. While brains can be manually dissected, for more efficient throughput and to be able to sort enough GFP+ nuclei from populations of few neurons, we used undissected heads for nuclei isolation. We homogenized flies with a Dounce homogenizer, with a filtration step between homogenization with the “A” and “B” pestles. The first step disrupts the fly exoskeleton, as shown by empty fly heads in the crude homogenate, indicating that the tissue inside the head is pushed out of the exoskeleton during homogenization. Passing the crude homogenate through a cell strainer eliminates much of the exoskeleton debris. Subsequent homogenization lyses the plasma membrane without disrupting the nuclear membrane (Corces et al., 2017). These freezing and homogenizing steps allowed us to recover up to 80% of GFP+ nuclei when ubiquitously expressing nuclear GFP. Thus, our physical extraction protocol allows for efficient and sensitive recovery of GFP-labelled nuclei from adult *Drosophila* heads.

An advantage of using *Drosophila* for ATAC-seq studies is the ability to genetically label nuclei of interest using the *Gal4/UAS* system. Thousands of Gal4 lines labeling distinct neurons are available (Jenett et al., 2012), including ones specific to neurotransmitters (Deng et al., 2019), allowing for the specific, reproducible interrogation of only a defined subset of neurons. As mentioned above, we were able to recover up to 80% of ubiquitous GFP+ nuclei at the low stringency gate, indicating good sensitivity. To determine the specificity, we sorted nuclei from GFP-negative flies and recovered 0.1% and 0.025% false positive nuclei at the low- and high-stringency gates. While these seem to be acceptable numbers for GFP+ specificity, they may present a problem when sorting from Gal4 lines with sparse expression. Sorting at the high-stringency gate will produce fewer false positives but will also reduce the sensitivity for collecting true positives. Possible solutions to the issue of insufficient GFP+ nuclei recovered include increasing the numbers of fly heads and increasing the GFP signal. We found that increasing the number of *Gal4* and *UAS-GFP-nls* transgenes increased the number of recovered GFP+ nuclei. The versatile *Gal4/UAS* system therefore allows for some flexibility to be able to sort ≥50,000 nuclei for ATAC-seq library preparation. We recommend that specific *Gal4* and *UAS-GFP-nls* combinations be tested by sorting nuclei to get information about the recovery of GFP+ nuclei prior to performing actual experiments consisting of library preparation and deep sequencing of experimental and control flies.

The *Drosophila* genome is ∼10-fold smaller than mammalian genomes, so enzyme-mediated steps during ATAC-seq library preparation may be affected by the amount of available DNA. Tn5 tagmentation in mammalian samples is generally performed using undiluted Tn5 enzyme included in a library preparation kit (1X) for 30 min at 37 °C (Buenrostro et al., 2015). Because the amount of accessible chromatin in *Drosophila* nuclei may be lower than in mammalian nuclei, we investigated the effects of changing the parameters of the enzymatic reactions (Tn5 tagmentation and PCR amplification) on fragment distribution and library quality. We observed that increasing or decreasing the Tn5 concentration reduced library complexity. Previous studies showed that decreasing the Tn5 concentration in mouse embryonic stem cells only decreases tagmentation efficiency when diluted to 10 nM or lower (Corces et al., 2017). Notably, this experiment used homemade Tn5 (Picelli et al., 2014), which may have different kinetics or concentrations than commercially-available Tn5 enzyme. Increasing the tagmentation time decreased library complexity, while decreasing the tagmentation time increased the number of mid-length fragments but did not improve nucleosomal periodicity. We also examined the number of PCR cycles needed to amplify ATAC-seq libraries before sequencing. Protocols for generating ATAC-seq libraries from mammalian tissues call for amplification to one-third the total fluorescence in a qPCR side reaction (Buenrostro et al., 2015). Because of the smaller genome size in *Drosophila*, amplifying ATAC-seq libraries to one-third of the total qPCR fluorescence may introduce GC and size bias. Indeed, amplifying our libraries to one-third qPCR fluorescence introduced a PCR bubble, indicating that the available primers were depleted during amplification (Kanagawa, 2003), causing amplification to occur using annealed adaptor sequences and increasing the mid-size fragments in the library. While these libraries may be sequenceable after one cycle of reconditioning PCR (Thompson et al., 2002), libraries with PCR bubbles should be quality-checked to verify that the PCR bubble was resolved before sequencing. Amplifying ATAC-seq libraries to more than 33% total qPCR fluorescence biases the library toward smaller fragment sizes and reduces library complexity. Based on these results, we recommend transposition with 1X Tn5 for 23 min, followed by amplification of the ATAC-seq libraries to 25% qPCR fluorescence, which preserves library complexity and does not introduce GC or size bias.

Using these optimized conditions, we generated ATAC-seq libraries from 65,000 nuclei isolated from *Drosophila* dopaminergic and GABAergic neurons. These libraries were of high quality, with the expected nucleosomal banding patterns and fragment size distributions. Additionally, these libraries had high alignment rates to the genome (92.4-96.5%), further confirming their quality. Therefore, we propose that ATAC-seq libraries from *Drosophila* tissues should use the following parameters (Fig. 8): freezing at -80 °C for no more than 5 min, at least three washes, 23-min tagmentation time with 1X Tn5, and amplification to 25% qPCR fluorescence. Our protocol harnesses the powerful genetic tools available in *Drosophila* to achieve excellent cell-type specificity followed by ATAC-seq, which is used to interrogate the chromatin landscape with unprecedented resolution. Thus, our protocol will enable detailed investigations of the role of chromatin-mediated gene regulation in normal and pathological states.

**Figure 8:**
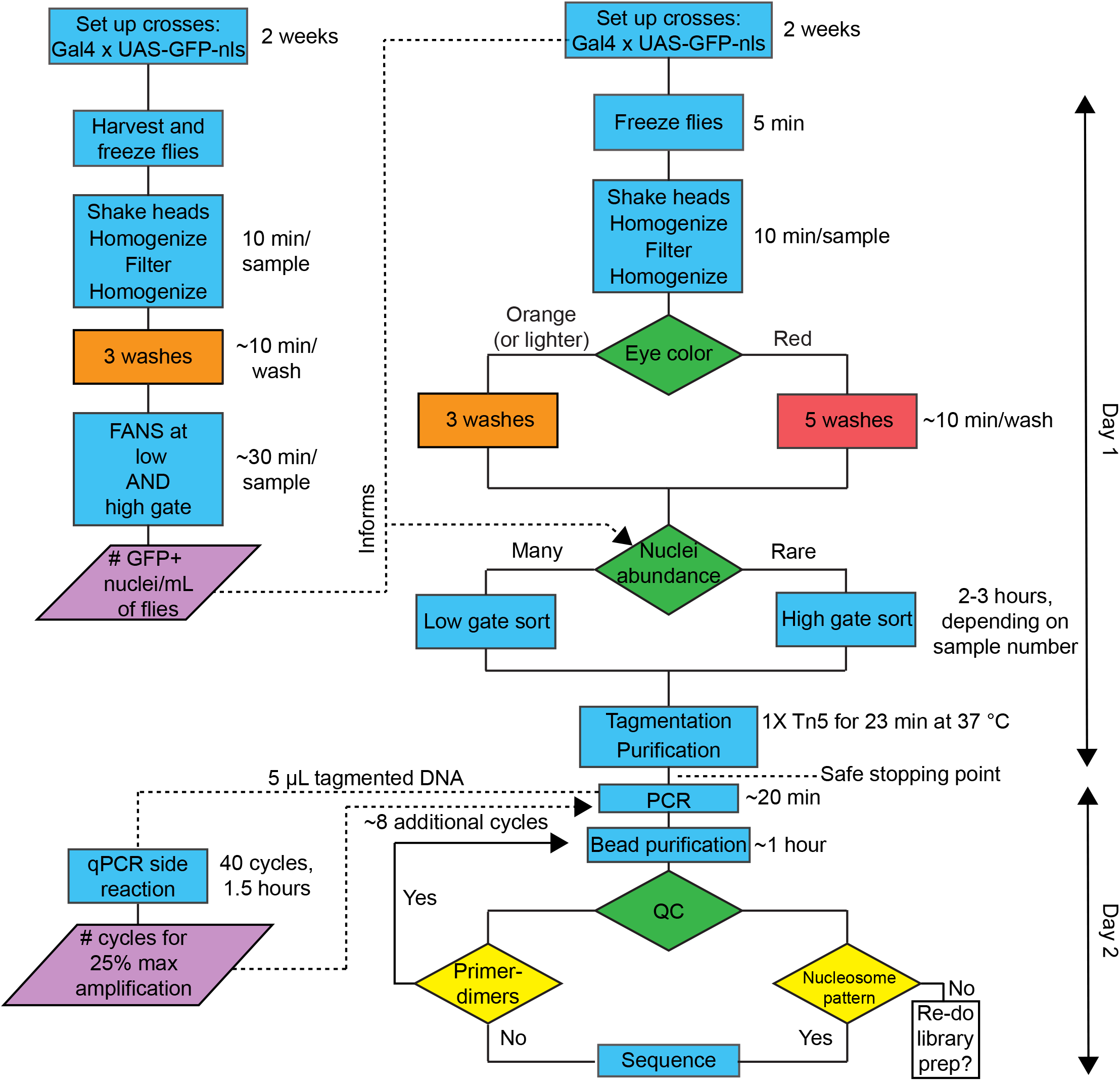
Flow chart of optimized ATAC-seq protocol. Suggested workflow, which we routinely use to generate ATAC-seq libraries from adult *Drosophila*. The libraries prepared using the workflow conform to ENCODE guidelines, with typical markers indicating a high-quality library, including nucleosomal banding pattern, high genome alignment rates, and high fraction of reads in peaks (FRiP). In the workflow, blue boxes indicate procedural steps, green diamonds indicate decision points, and yellow diamonds indicate decision points based on QC metrics. On the right of each procedural box is shown the approximate time required for each step.

## ACKNOWLEDGMENTS

This work was supported by the University of Utah Flow Cytometry Facility, the University of Utah Genomics Core Facility, the High Throughput Sequencing Core at the Huntsman Cancer Institute, and the National Cancer Institute through Award Number 5P30CA042014-24. The content is solely the responsibility of the authors and does not necessarily represent the official views of the National Institutes of Health.

## AUTHOR CONTRIBUTIONS

CBM performed ATAC-seq experiments, analyzed data; and wrote the manuscript; MAP performed nuclei isolation, sorting, and optimization experiments; ABM analyzed data; ARR reviewed and edited the manuscript; AR supervised the project and revised and edited the manuscript. All authors reviewed the manuscript.

## DATA AVAILABILITY

The datasets generated by this study are available upon reasonable request.

